# Structural and biochemical characterisation of an iterative GCN5-related *N*-acetyltransferase required for fungal siderophore tailoring

**DOI:** 10.64898/2026.09.24.754016

**Authors:** Joseph D. Newman, Y. T. Candace Ho, Christopher D. Fage, Günther Winkelmann, Lona M. Alkhalaf, Józef R. Lewandowski, Fabrizio Alberti, Matthew Jenner

## Abstract

Siderophore-mediated iron acquisition is essential for fungal survival, particularly under iron-limiting conditions. In *Aspergillus fumigatus*, SidG, a member of the GCN5-related *N*-acetyltransferase (GNAT) superfamily, catalyses the final step in the biosynthesis of the extracellular siderophore triacetylfusarinine C (TAFC) through sequential acetylation of the precursor fusarinine C (FsC). However, the timing, catalytic mechanism, and functional significance of this modification are not fully understood. Here, we reconstituted SidG activity *in vitro* and combined native mass spectrometry, X-ray crystallography, molecular dynamics simulations, and site-directed mutagenesis to investigate its catalytic properties. Our analyses demonstrate that SidG selectively binds acetyl-CoA from the cellular milieu and iteratively acetylates the FsC scaffold prior to iron chelation. Structural, biochemical, and molecular dynamics analyses support a direct transfer mechanism, identify key catalytic residues, and demonstrate the strict selectivity of SidG for short-chain acyl-CoA donors. Together, these findings establish the molecular basis for SidG-dependent siderophore tailoring and expand our understanding of GNAT-catalysed transformations in fungal natural product biosynthesis.

## INTRODUCTION

Iron is an essential nutrient for microbial life, underpinning numerous fundamental cellular processes.^1,2^ However, despite its natural abundance, bioavailable iron is often scarce. In aerobic environments it predominantly exists as Fe^3+^, which readily forms insoluble ferric hydroxide species, rendering it largely inaccessible to microorganisms.^3^ To overcome this limitation, bacteria and fungi have evolved diverse strategies for iron acquisition and storage.^4^ Among these is the biosynthesis of siderophores; low-molecular-weight, high-affinity metal chelators that solubilise and sequester ferric iron.^5,6^

Fungal siderophores typically contain hydroxamate functional groups, three of which coordinate Fe³⁺ in a hexadentate arrangement.^7^ A prominent example is triacetylfusarinine C (TAFC), produced by *Aspergillus fumigatus* and other fungi. The biosynthesis of TAFC begins with the nonribosomal peptide synthetase (NRPS), SidD, which assembles three *N*^5^-*cis*-anhydromevalonyl-*N*^5^-hydroxy-L-ornithine (*cis*-AMHO) residues into the depsipeptide fusarinine C (FsC).^8^ The newly formed FsC then undergoes acetylation of its three amine groups by SidG, a GCN5-related *N*-acetyltransferase (GNAT) superfamily enzyme (**Fig. 1**), although the functional significance of this acetylation is currently unclear. Deletion of the *sidG* gene results in a 10-fold accumulation of the FsC precursor without affecting virulence or growth under iron-limited conditions, suggesting that elevated FsC levels can compensate for the absence of TAFC.^9^ Interestingly, two distinct MFS transporters mediate the uptake of these siderophores from the environment: MirB imports TAFC, whereas MirD imports FsC (**Fig. 1**).^10,11^ Similarly, the hydrolases required for intracellular release of iron also show selectivity: EstB recognises TAFC, whereas SidJ processes FsC (**Fig. 1**).^12–14^ These observations suggest that acetylation occurs after assembly of the FsC scaffold. Indeed, acetylation of the individual *cis*-AMHO units seems unlikely, as recognition by NRPS adenylation domains typically requires a positively-charged amine functionality, which forms an important salt bridge with a conserved Asp residue.^15–17^ Likewise, modification of Fe^3+^–FsC also seems improbable, as Fe^3+^ coordination would likely restrict the conformational space that the macrocycle can explore. Whilst these considerations point to acetylation of the newly formed FsC prior to Fe^3+^ chelation, this remains experimentally unverified.

**Figure 1.**
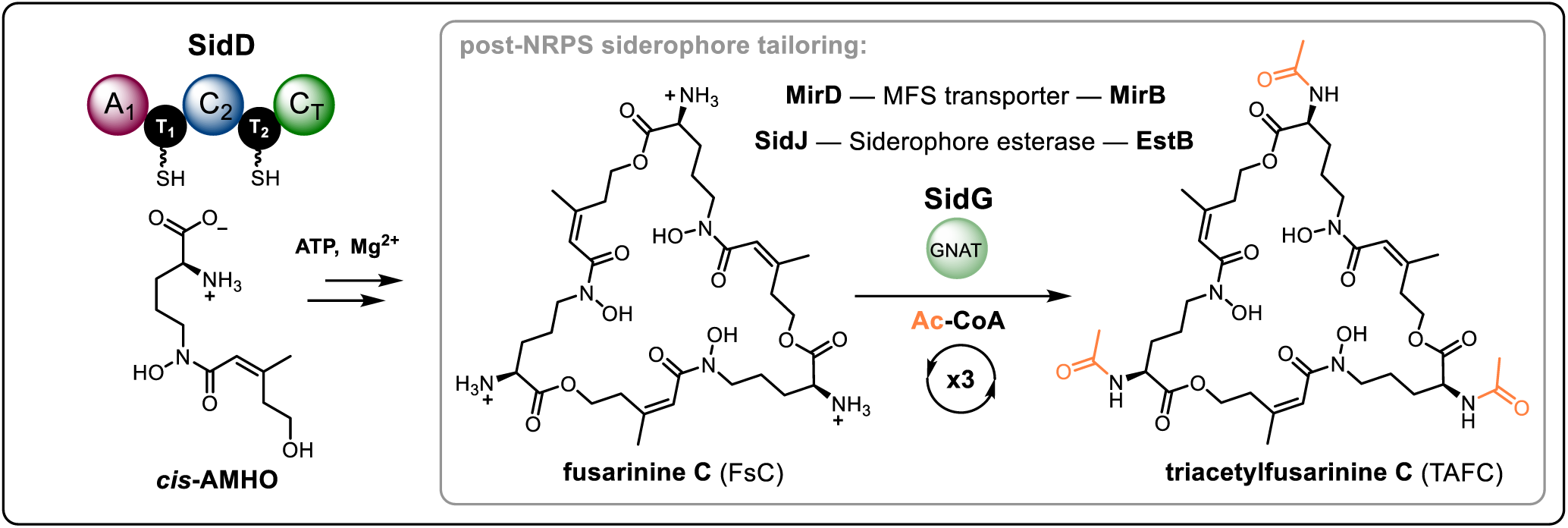
SidG catalyses acetylation of fusarinine. **C.** Schematic showing the biosynthesis of triacetylfusarinine (TAFC). Three *cis*-AMHO units are condensed by the SidD NRPS to yield fusarinine C (FsC), which is subsequently acetylated by SidG to yield TAFC. The acetyl groups are highlighted in orange and the siderophore-specific transporters and esterases are shown. NRPS domains: adenylation (A), thiolation (T), condensation (C) and terminal condensation (CT). A wavy bond to S indicates the phosphopantetheine cofactor that covalently tethers the biosynthetic intermediate to T domains.

The mechanism and substrate recognition of SidG are also of interest. GNAT enzymes typically catalyse direct acetyl transfer from acetyl-CoA to an acceptor substrate via formation of a ternary complex, with a conserved aspartate or glutamate residue often acting as a general base to activate a primary amine nucleophile.^18^ However, mechanistic diversity has been reported within the GNAT superfamily, including formation of a covalent acyl-enzyme intermediate in hybrid ping-pong mechanisms.^19,20^ While the canonical mechanism is well established, it remains unclear whether the fungal enzyme, SidG, follows the same catalytic strategy. An additional consideration is that SidG must function iteratively during TAFC biosynthesis, acting on mono- and di-acetylated intermediates as well as FsC itself. The progressive neutralisation of amino groups alters the charge and chemical properties of the substrate at each step, raising questions as to how SidG accommodates and correctly orients these distinct species for catalysis.

Here, we reconstitute SidG activity *in vitro* and combine structural, biochemical, and mass spectrometric approaches to investigate its role in FsC tailoring, its catalytic mechanism, and its substrate scope. Native mass spectrometry was used to probe substrate binding and monitor acetylation, and X-ray crystallography was employed to determine the structure of SidG in complex with acetyl-CoA. Molecular dynamics simulations and site-directed mutagenesis were further used to explore the catalytic mechanism. Finally, substrate scope analyses were undertaken to assess the ability of SidG to utilise alternative acyl-CoA donors and generate corresponding acylated derivatives of FsC.

## RESULTS AND DISCUSSION

### Production and mass spectrometric analysis of recombinant SidG

To begin investigating acetylation of FsC, SidG from *A. fumigatus* A293 was overproduced in *Escherichia* coli as an N-terminal His^8^-tagged fusion protein and subsequently purified to homogeneity using immobilized metal-ion affinity chromatography. Visualisation by SDS-PAGE yielded a single major band (**Supplementary Fig. 1a**), however, analysis by UPLC-ESI-Q-TOF-MS revealed a range of degradation products (**Supplementary Fig. 1b**). The major species corresponded to a loss of ∼2.7 kDa from the expected mass, with C-terminal truncation being the most likely explanation, as deletion of these residues from the predicted sequence resulted in masses matching the observed values. Repeating the purification in the presence of protease inhibitors yielded the same degree of degradation. An AlphaFold model of SidG predicted a highly disordered C-terminal region (Pro195 – Pro235), which did not associate with the core GNAT fold (**Supplementary Fig. 1c**). Suspecting that the C-terminal region was subject to degradation in *E. coli*, a truncated SidG (Δ199–235) construct was generated. Following purification and removal of the N-terminal His-tag, the protein yielded a single intact species by UPLC-ESI-Q-TOF-MS analysis, with no evidence of additional degradation (**Supplementary Fig. 1d** and **e**). This truncated construct is referred to as SidG from this point forward.

Native mass spectrometry analysis of purified SidG exhibited two charge-state series in the range of 9^+^ to 11^+^. The first corresponded to a monomeric SidG species, consistent with analytical size-exclusion chromatography measurements (**Fig. 2a** and **Supplementary Fig. 2**). However, the second series corresponded to ions that were +809 Da in mass, indicating non-covalently bound acetyl-CoA (**Fig. 2a**). Indeed, isolation of the corresponding 10^+^ charge state (*m/z* = 2379.4) followed by collision-induced activation of the complex yielded a diagnostic fragment ion for acetyl-CoA (*m/z* = 303.1), arising from cleavage of the labile phosphoester linkage (**Fig. 2b**). Co-purification of SidG with acetyl-CoA suggests a high affinity for the endogenous cofactor present in the cell. Interestingly, the bound ligand appears to be exclusively acetyl-CoA rather than CoA, suggesting interactions with the acetyl moiety, in addition to the CoA scaffold, favour the acetylated form.

**Figure 2.**
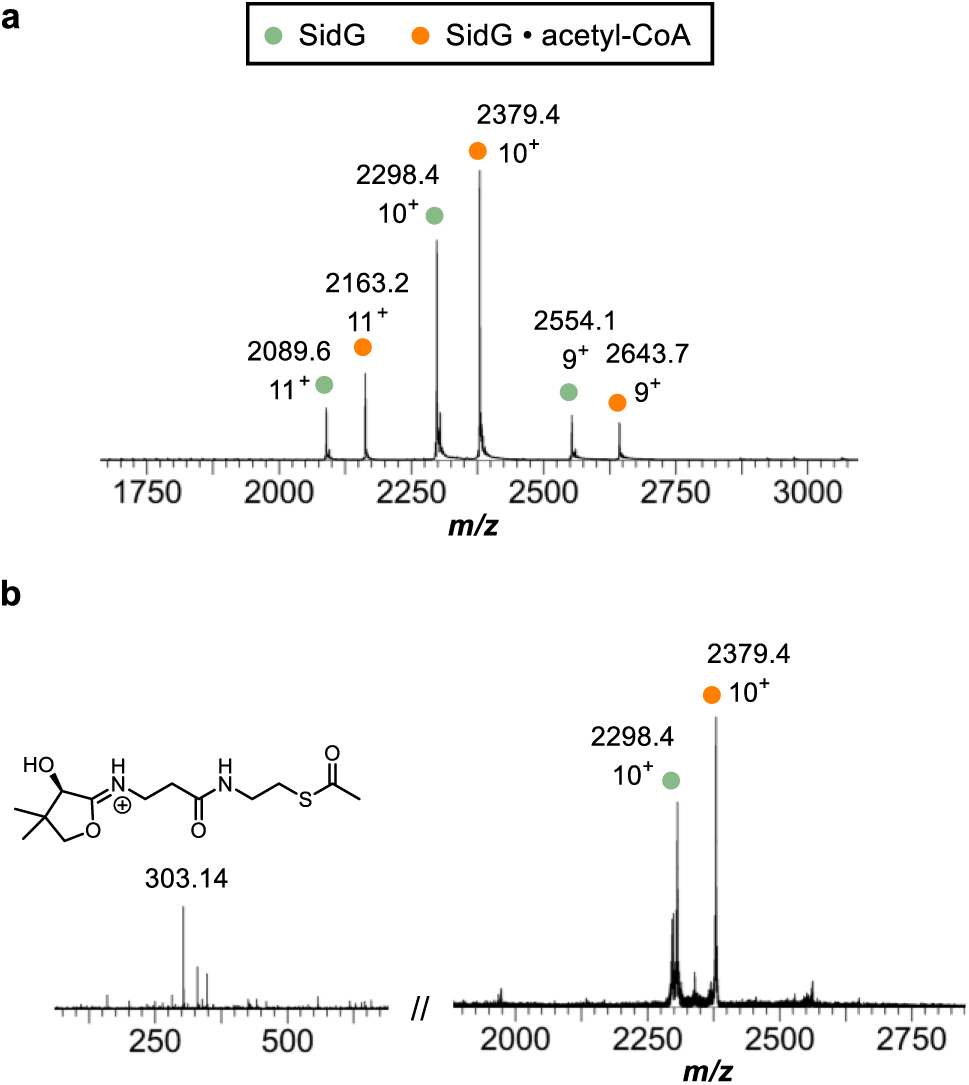
Recombinant SidG purifies with acetyl-CoA bound. (a) Native mass spectrum of SidG sprayed from 50 mM NH_4_OAc showing two charge-state distributions from 9^+^ to 11^+^. The green dots indicate the SidG protein alone, and the orange dots the SidG • acetyl-CoA complex. (b) Mass spectrum following isolation and collision-induced dissociation of the 10^+^ SidG • acetyl-CoA species (*m/z* = 2379.4), which yields a diagnostic fragment ion from acetyl-CoA (chemical structure depicted).

### Reconstitution of SidG-catalysed acetylation of FsC

We next sought to reconstitute the acetylation reaction *in vitro* using purified FsC as the substrate and acetyl-CoA as the acyl donor. Incubation of SidG with FsC and acetyl-CoA resulted in the formation of TAFC after 30 seconds, with mono-acetylated (MAFC) and di-acetylated (DAFC) species detectable after 10 seconds (**Fig. 3a**). Indeed, a time-course analysis of the reaction indicated completion within 1 minute (**Supplementary Fig. 3**). In contrast, an enzyme-free control failed to produce any acetylated products, demonstrating that, within the timeframe of this assay, non-enzymatic acyl transfer between acetyl-CoA and the amino groups of FsC does not occur (**Fig. 3a**). We were also able to visualise the reaction using native protein MS. Noting that SidG co-purifies with acetyl-CoA in ∼50 % occupancy (**Fig. 2a**), addition of FsC alone immediately prior to MS infusion allowed observation of a new charge-state series (9^+^ to 11^+^) corresponding to a set of species with increased masses of +768, +810, and +852 Da relative to the SidG • acetyl-CoA complex (**Fig. 3b**). These mass shifts are consistent with the formation of ternary complexes of SidG • acetyl-CoA • MAFC / DAFC / TAFC, respectively. To verify the identity of these species, we subjected the 10^+^ charge state to collisional activation with a wide isolation window to capture all peaks. This resulted in the release of an ion at *m/z* = 853.4, consistent with the mass of TAFC, confirming its formation within the complex (**Fig. 3b**). The absence of MAFC and DAFC release upon collisional activation is likely due to strong ionic interactions between the –NH_3_^+^ groups of these substrates and negatively-charged residues in the SidG active site (*vide infra*), which are abolished in the TAFC form. Additionally, the relative signal intensities shifted from SidG • acetyl-CoA toward SidG alone, and free CoA was detected (*m/z* = 768.1), further supporting catalytic turnover and release of products.

**Figure 3.**
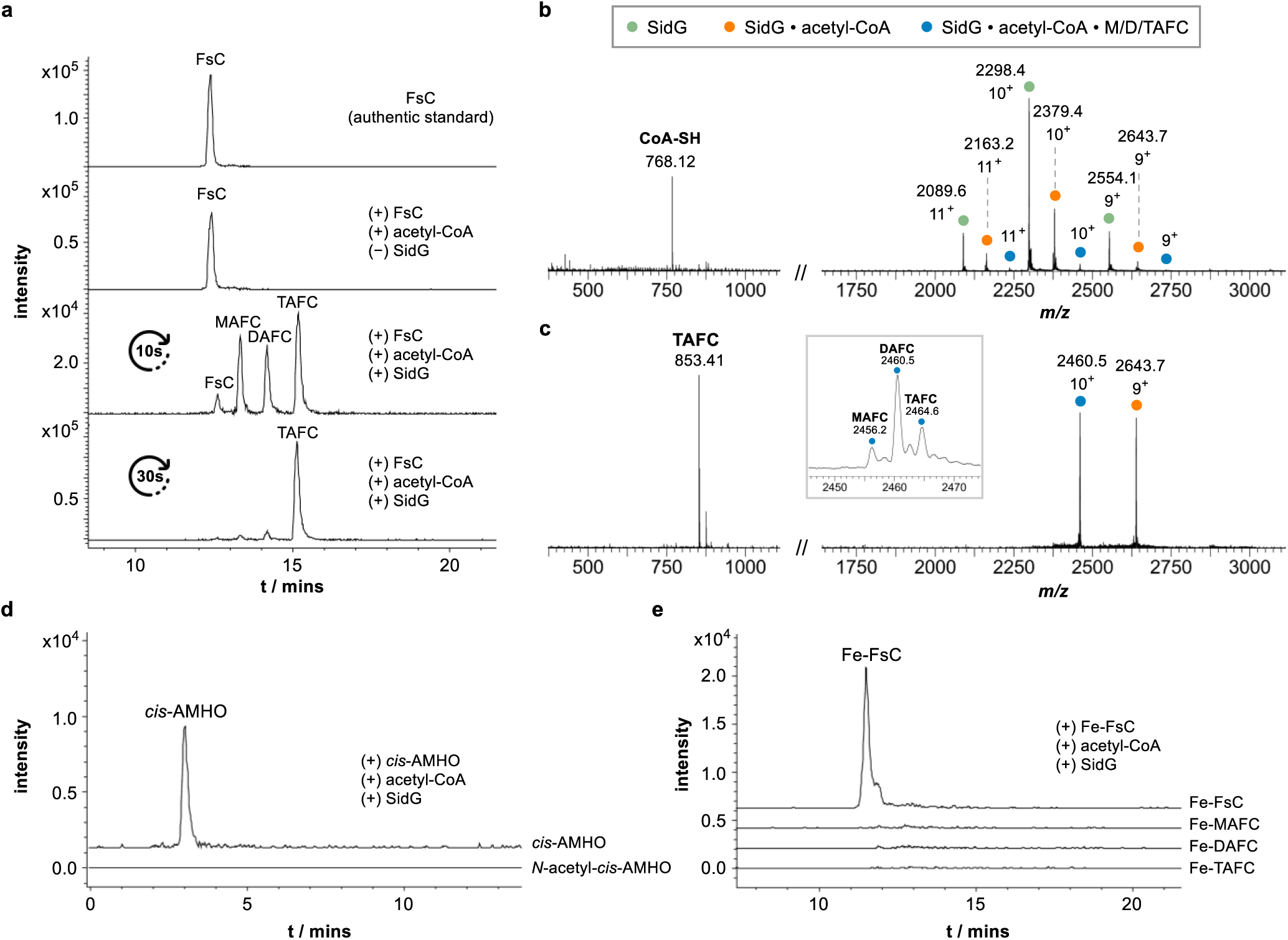
*In vitro* reconstitution of SidG-catalysed acetylation activity. (a) LC-MS chromatograms showing progression of SidG-catalysed acetylation of FsC. Quenching the reaction after 10s enabled observation of mono-, di-, and tri-acetylated species before progression to the completely tri-acetylated product. Each chromatogram is a combined extracted ion chromatogram of FsC (*m/z* = 727.39, 364.20 ± 0.01); MAFC (*m/z* = 769.40, 385.20 ± 0.01); DAFC (*m/z* = 811.41, 406.21 ± 0.01); and TAFC (*m/z* = 853.42, 427.21 ± 0.01). (b). Native mass spectrum of SidG sprayed from 50 mM NH_4_OAc following incubation with FsC. Compared to Fig. 2b, a reduction in the SidG • acetyl-CoA complex (orange dots) is observed, and the emergence of a charge-state series corresponding to the SidG • CoA • MAFC / DAFC / TAFC complexes (blue dots, see inset). (c) Mass spectrum following isolation and collision-induced dissociation of the 10^+^ SidG • CoA • M/D/TAFC species (*m/z* = 2456 – 2464), which yields a species at *m/z* = 853.41, corresponding to the [M+H]^+^ for TAFC. MAFC / DAFC were not observed. (d) LC-MS chromatograms showing that *cis*-AMHO (*m/z* = 261.14 ± 0.01) is not acetylated by SidG to form *N*-acetyl-*cis*-AMHO (*m/z* = 303.16 ± 0.01). (e) LC-MS chromatograms showing that Fe-FsC (*m/z* = 780.30, 390.65 ± 0.01) is not acetylated by SidG to form the corresponding Fe-MAFC (*m/z* = 822.31, 411.66 ± 0.01); Fe-DAFC (*m/z* = 864.32, 432.66 ± 0.01); and Fe-TAFC (*m/z* = 906.33, 453.67 ± 0.01) species.

SidG was also tested for its ability to acetylate two related compounds: *cis*-AMHO (i.e. the monomeric unit for FsC assembly) and Fe^3+^–FsC (i.e. the product of iron capture prior to acetylation), and no acetylation was detected for either substrate under the assay conditions after 1 hour (**Fig. 3d** and **e**). These results indicate that SidG does not recognise the smaller *cis*-AMHO substrate, implying that only non-acetylated monomers are incorporated during FsC assembly, which is in agreement with previous work.^8,17^ Rather, it exclusively acts directly after FsC assembly, acetylating the free amine groups before the siderophore is secreted for iron scavenging. Furthermore, despite both FsC and TAFC being secreted by *A. fumigatus* A293 for iron acquisition, these data suggest that once FsC has bound Fe^3+^ acetylation by SidG can no longer occur, with iron binding likely inducing an unfavorable conformation.

### Structural analysis of SidG and substrate binding pockets

To elucidate the catalytic mechanism and substrate-binding properties of SidG, we determined its crystal structure. Here, a preparation of SidG (10 mg/mL) was subjected to sparse-matrix crystal screening, which yielded diffraction-quality crystals. However, analysis of the diffraction datasets in the Phenix software package^21^ suggested the presence of twinning, which hindered structure determination. As previously noted, native protein MS analysis indicated that ∼50 % of the enzyme population was bound to acetyl-CoA (**Fig. 2a**), raising the possibility that the presence of both ligand-bound and -unbound states introduced heterogeneity that complicated structure determination. Whilst investigating active site mutations to probe the enzymatic mechanism (*vide infra*), we identified an S149A variant of SidG that exhibited a substantially higher proportion of acetyl-CoA-bound protein (∼90%) as determined by native protein MS (**Supplementary Fig. 4**). Although the reason for the enhanced incorporation is unclear, the increased occupancy facilitated subsequent crystallographic studies. Indeed, crystals obtained using the same conditions as for wild-type SidG yielded a native X-ray diffraction dataset to a resolution of 2.1 Å (PDB: 32FB).

The SidG structure exhibits a typical GNAT fold, consisting of five α-helices (α1 – α5) that surround a predominantly antiparallel β-sheet core, made of seven strands (β1 – β7). A β-bulge between β4 and β5 results in the characteristic splay in the sheet, providing access for acetyl-CoA (**Fig. 4a**).^18^ Electron density was missing in loop regions connecting α1 to α2 (M39 – G43) and β3 to β4 (V88 – R98), presumably due to the high conformational mobility of these regions. These missing residues were added using MODELLER to produce a contiguous polypeptide chain.^22^ Electron density was located between α3 / α4 and β4 / β5 into which acetyl-CoA could be modelled, and in agreement with our native protein MS measurements (**Fig. 4b** and **Supplementary Fig. 4 and 5**). Here, the 3ʹ-phosphate of the acetyl-CoA substrate makes a stabilising contact with the K119 side chain, and the 5ʹ-diphosphate with the backbone of α3, whilst the carbonyl of the β-alanine region interacts with the N146 side chain. Interestingly, the carbonyl of the acetyl group is interacting with the backbone amide of Phe106, rather than occupying the proposed oxyanion hole formed by N146 and T142 (**Fig. 4b**).^19^

**Figure 4.**
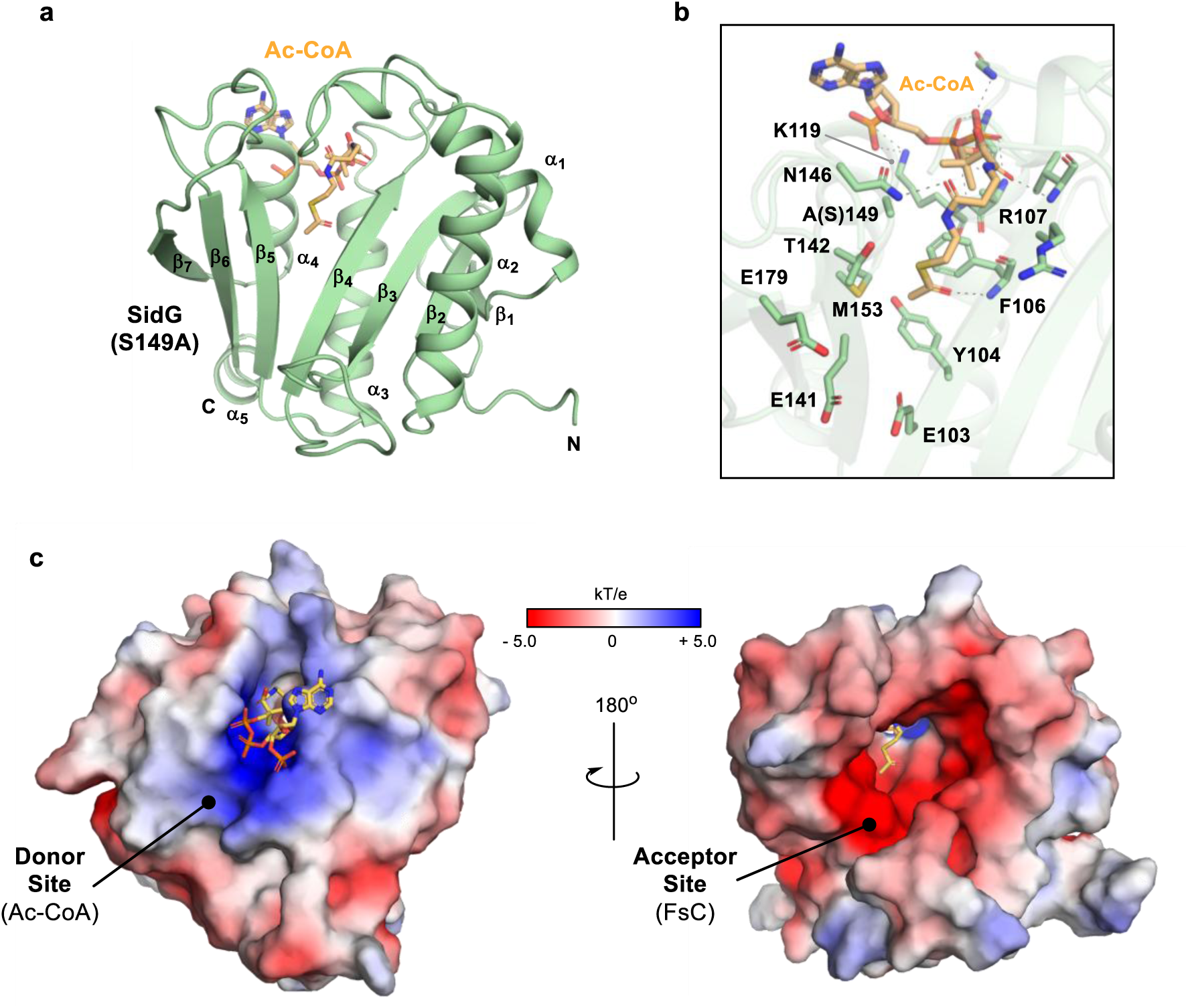
Structure of the SidG (S149A) • Ac-CoA complex and binding site analysis. (a) Crystal structure of the SidG (S149A) acetyl-CoA complex (PDB: 32FB). The protein structure is shown in cartoon with secondary structure elements labelled, and acetyl-CoA as sticks with orange C atoms. Loop regions connecting α1 to α2, β3 to β4, and β6 to β7 were added using MODELLER. (b) Active site of SidG showing residues proximal to the acetyl-CoA as sticks, with contacts to the acetyl-CoA ligand shown as black dashed lines. (c) Structure of SidG displaying solution-phase surface electrostatics calculated using APBS. The donor site (acetyl-CoA) is positively charged to stabilise the negatively-charged phosphates. In contrast, the acceptor site (FsC) on the reverse side of the enzyme is highly negatively charged to promote binding of the –NH_3_^+^ groups of FsC.

**Figure 5.**
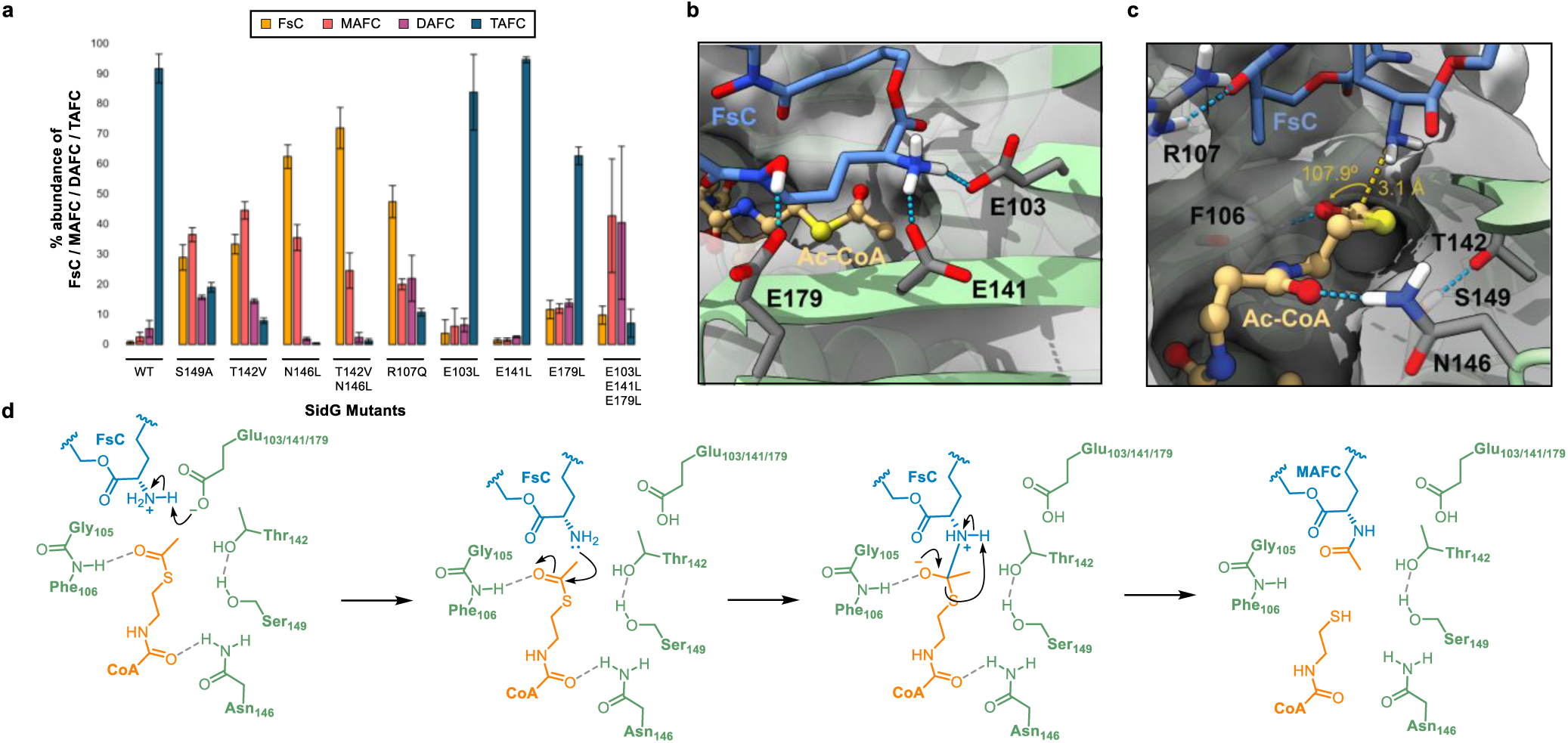
Mutagenesis and molecular dynamics suggest a direct transfer catalytic mechanism for SidG. (a) Bar chart showing SidG-catalysed formation of MAFC, DAFC and TAFC for active site variants compared to WT SidG activity. (b) Example frame from a cMD simulation of the SidG • acetyl-CoA • FsC ternary complex showing a catalytically competent geometry, in which the distance and angular requirements for nucleophilic attack by an FsC amine on the acetyl-CoA carbonyl are satisfied. (c) Representative frame from cMD simulations of the SidG • acetyl-CoA • FsC ternary complex showing key substrate-binding residues. The cluster of glutamate residues (E103, E141 and E179) that interact with the amino groups of FsC are highlighted, alongside the arginine residue (R107) that interacts with the hydroxamate functionality of FsC regularly throughout the simulations. (d) Proposed direct transfer mechanism of SidG for a single acetylation event of FsC to form MAFC.

Examination of the overall architecture and electrostatic surface reveals that the CoA-binding tunnel, corresponding to the donor site, is both highly constricted and predominantly positively charged. The narrow geometry may function to exclude branched acyl-CoA species, while the positive electrostatic environment likely stabilises the negatively charged CoA moiety during binding (**Fig. 4c**). In contrast, the acceptor site, located on the opposite face of the enzyme, forms a large negatively charged cavity capable of accommodating FsC. This large active site cavity contains several areas of electron density that can be reasonably modelled as glycerol units in the crystal structure. At physiological pH, the substrate amine groups are expected to be predominantly in the protonated (NH3^+^) form. The negatively-charged cavity, largely formed by Glu residues, likely promotes substrate binding and may facilitate amine deprotonation during catalysis (*vide infra*). The expansive nature of the cavity likely contributes the required conformational flexibility to accommodate changes in the substrate as it is converted from FsC to MAFC and subsequently to DAFC (**Fig. 4c**).

### Mutagenesis and molecular dynamics simulations to probe mechanism and substrate binding

Our high-resolution structure of the SidG • acetyl-CoA complex provided a framework for rational mutagenesis and molecular dynamics simulations aimed at probing the catalytic mechanism and substrate-binding capabilities of SidG. The closet characterised homologue to SidG is the PA3944 GNAT enzyme (25.4% identity to SidG across 224 residues), which is responsible for acetylation of a single diaminobutyric acid residue in polymyxin B and E. Based on kinetic analyses and docking simulations, this enzyme is proposed to act via a hybrid ping-pong mechanism in which both direct acetyl transfer and an acyl-enzyme intermediate pathway may occur.^19,20,23^ Here, a nucleophilic Ser residue was the proposed site of acyl chain attachment, with a putative oxyanion hole formed by Thr and Asn side chains. These residues are conserved in SidG, corresponding to S149, T142 and N146, respectively (**Supplementary Fig. 6**). Although SidG purified with Ac-CoA bound (**Fig. 2a**), analysis by intact denatured protein MS did not reveal a mass shift corresponding to acetylation (+ 42 Da), even under conditions of excess acetyl-CoA (**Supplementary Fig. 7**). This observation is consistent with our structural data, which show the acetyl group positioned in a manner not obviously conducive to nucleophilic attack by S149: the carbonyl of the acetyl group interacts with the backbone NH of F106, instead of occupying an oxyanion-stabilising environment (putatively involving T142 and N146) (**Fig. 4b**).

Mutagenesis data further corroborate a mechanism that does not rely on a covalent intermediate for SidG. The S149A mutation did not abolish enzymatic activity, instead resulting in a reduction in TAFC formation compared to wild type, with MAFC and DAFC intermediates still detected (**Fig. 5a**). A similar, mild effect was observed for the T142V variant. These results suggest that neither residue is essential for catalysis. In contrast, an N146L mutation led to a pronounced decrease in TAFC production, with predominantly MAFC and only trace amounts of DAFC observed. This effect was slightly exacerbated in the N146L, T142V double mutant (**Fig. 5a**). Structural analysis reveals that the N146 side chain forms a hydrogen bond with the carbonyl group of the β-alanine moiety of coenzyme A (**Fig. 4b**), and disruption of this interaction likely impairs proper substrate positioning, thereby reducing catalytic efficiency. To explore whether acetyl-CoA could adopt a conformation compatible with acylation of Ser149, the SidG • acetyl-CoA complex was subjected to a series of 300 ns classical molecular dynamics (cMD) simulations. Although a small subset of frames from two out of three independent cMD simulations sampled near-attack conformations (Ser149 Oγ→C1 ≤ 3.5 Å; Oγ–C1–O = 100-110°), none exhibited insertion of the carbonyl oxygen into the putative oxyanion hole formed by T142 and N146; only infrequent hydrogen bonding to the T142 side chain was observed. (**Supplementary Fig. 8**). Although an oxyanion hole is not essential for acyl transfer chemistry, it is often required to facilitate nucleophilic attack by enhancing carbonyl polarisation and stabilising the resulting tetrahedral intermediate. The absence of interactions with the proposed oxyanion hole therefore limits support for efficient Ser acylation under the simulated conditions.

A direct-transfer mechanism requires formation of a ternary SidG • acetyl-CoA • FsC/MAFC/DAFC complex, which was observed for both MAFC and DAFC by native mass spectrometry (**Fig. 3b** and **3c**). To investigate this mechanism, we performed a series of 300 ns cMD simulations of the SidG • acetyl-CoA • FsC complex, in which all three amino groups of FsC were protonated (3 × –NH_3_^+^). As starting points, we used two models predicted by Protenix^24^, which differed in the orientation of FsC relative to acetyl-CoA. In all simulations, none of the NH_3_^+^ groups approached the acetyl-CoA thioester carbonyl close enough (< 4 Å) to permit nucleophilic attack. However, the amino group nearest to acetyl-CoA frequently formed interactions with the side chains of E103, E141, and E179 (**Supplementary Fig. 9**). Given the potential for each of these glutamate residues to act as a general base to deprotonate the FsC amino groups, we performed additional 300 ns cMD simulations initiated from frames in which an –NH_3_^+^ group was simultaneously hydrogen bonded to two glutamates (E179/E141 or E141/E103). Assuming deprotonation by one of these residues, the interacting amino group was converted to NH_2_ prior to simulation. In all instances, the neutral –NH_2_ group sampled positions much nearer to the acetyl-CoA carbonyl than its protonated counterpart, and in several simulation replicates adopted geometries consistent with a near-attack conformation relative to the acetyl-CoA thioester carbonyl (FsC NH_2_ → C1 ≤ 3.5 Å; N–C1–O = 100–110°), with the acetyl-CoA carbonyl oxygen stabilised by hydrogen bonding to the backbone amide of F106 in all cases (**Fig. 5c** and **Supplementary Fig. 10**). In several additional trajectories, near-attack conformations were only observed after extending the simulations by a further 300 ns.

Taken together, near-attack conformations were observed following putative deprotonation by E103, E141, and E179. The ability of each residue to support formation of reactive geometries suggests a degree of functional redundancy within the active site, whereby loss of any single glutamate can be compensated by the remaining residues. This is consistent with the mutagenesis data, in which individual substitutions had only modest effects on activity, whereas simultaneous mutation of all three residues substantially impaired catalysis (**Fig. 5a**). Interestingly, whilst E103 is conserved between SidG and the PA3944 GNAT, E141 and E179 are replaced by Phe and His residues, respectively, suggesting that these residues represent an adaptive feature of SidG that promotes productive FsC binding (**Supplementary Fig. 6**).

Throughout these simulations, acetyl-CoA and FsC engaged in a dynamic network of interactions with several active site residues. The N146 side chain interacted predominantly with the carbonyl of the β-alanine region of CoA, while the T142 and S149 side chains alternated between hydrogen bonding to one another (an interaction that stabilises the relative positions of the helix and β-strand, thereby influencing acetyl-CoA and FsC orientation in the active site), and transiently interacting directly with acetyl-CoA or FsC. These observations are consistent with the reduced, but not abolished, catalytic activity observed upon mutation of these residues (**Fig. 5a**). Notably, the R107 side chain was observed to form persistent interactions with the hydroxamate region of FsC, helping to position the macrocycle within the active site. Consistent with an important role in substrate recognition and orientation, similar to that observed for PA3944 GNAT^20^, an R107Q mutant resulted in a substantial reduction in catalytic activity (**Fig. 5a** and **Supplementary Fig. 11**). Importantly, this interaction is likely lost in the ferric complex, as the hydroxamate oxygens are engaged in Fe^3+^ coordination and are therefore unavailable for interaction with R107. Combined with the conformational constraints imposed by metal chelation, this loss of substrate positioning likely contributes to the inability of SidG to acetylate Fe^3+^–FsC (**Fig. 3e**).

The electrostatic interactions between positively charged NH_3_^+^ groups and glutamate residues lining the binding pocket are expected to be retained for FsC, MAFC and DAFC, but progressively diminished during the iterative cycle of catalysis, thereby facilitating product release. Consistent with this model, native mass spectrometry revealed that only TAFC could be readily dissociated from SidG under collisional activation conditions, whereas complexes containing earlier intermediates remained resistant to dissociation, indicative of stronger residual binding interactions (**Fig. 3** and **3c**). To investigate how the interaction network evolves following acetylation of FsC, we performed 150 ns accelerated molecular dynamics (aMD) simulations of the SidG • acetyl-CoA • MAFC complex. In all three independent aMD replicates, which were used to sample a broader region of conformational space than cMD simulations of equivalent duration, the acetylated amine departed from the active site while a second NH_3_^+^ group approached. This process appeared to be guided by favourable interactions with negatively charged active-site residues (**Supplementary Movie 1**). In one of the simulations, acetyl-CoA was observed leaving the binding pocket, with both of the unacetylated NH_3_^+^ groups visiting the active site over the course of the simulation. In the other independent repeats, where acetyl-CoA remains in SidG, only one of the unacetylated NH_3_^+^ sites ends up in a similar position. Taken together, these findings support a mechanism in which precise alignment of acetyl-CoA, rather than formation of a covalent intermediate, is critical for efficient acetyl transfer in SidG (**Fig. 5d**).

### Insights into acyl-CoA chain length control

Our data demonstrate that SidG co-purifies with bound acetyl-CoA, with neither CoA-SH nor any alternative acyl-CoA species detected (**Fig. 2a**), suggesting a strong preference for a C_2_ acyl chain. To investigate whether SidG can utilise longer acyl-CoA substrates, we performed our standard assay using propionyl-, butyryl-, hexanoyl-, and octanoyl-CoA, and monitored formation of the corresponding acyl-FsC products. Propionyl-CoA was processed with an efficiency comparable to that of acetyl-CoA; however, product formation decreased markedly with butyryl-CoA, and no turnover was observed with either hexanoyl-or octanoyl-CoA (**Fig. 6a**). Although all non-native acyl-CoA substrates were supplied in excess, residual acetyl-CoA co-purified with SidG was also present in these assays, leading to low levels of MAFC / DAFC / TAFC formed during the assays. Frames from our MD simulations capturing catalytically competent conformations revealed the acyl moiety of acetyl-CoA is positioned within a shallow binding pocket that is optimally sized for a C_2_ chain (**Fig. 6b**). This architecture likely imposes steric constraints that disfavour accommodation of longer acyl chains, consistent with the observed substrate specificity. Although the substrate specificity of SidG could potentially be broadened through rational mutagenesis or directed evolution, the practical utility of such engineering is limited, as FsC derivatives bearing alternative acyl substituents are readily accessible through established synthetic procedures.^25^

**Figure 6.**
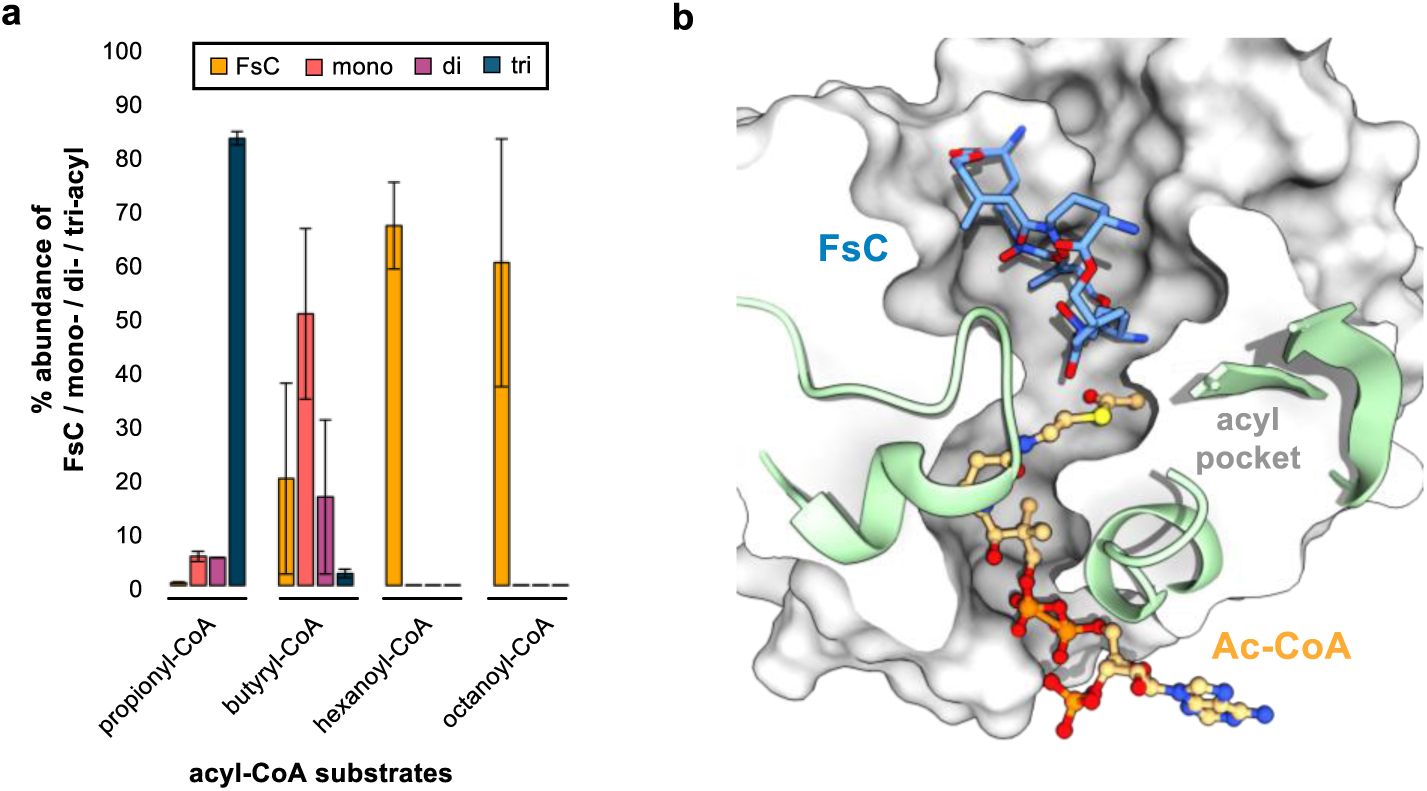
Structural basis for acyl chain-length selectivity in SidG. (a) Bar chart showing SidG-catalysed formation of mono- / di- / tri-acylated forms of FsC upon incubation with acyl-CoAs with increasing chain length. (b) Example frame from cMD simulations of the SidG • acetyl-CoA • FsC ternary complex in a catalytically competent geometry. The constrained acyl binding pocket limits the acyl chain length accepted by SidG.

## CONCLUSIONS

Our detailed study of SidG clarifies the order of events in TAFC biosynthesis, demonstrating that acetylation occurs after NRPS-mediated assembly of FsC and prior to ferric iron acquisition. Biosynthetic fidelity for TAFC is achieved through the strict selectivity of SidG for acetyl-CoA, which is conferred by a constricted acyl-binding pocket. The data support a direct acetyl transfer mechanism from acetyl-CoA within a ternary enzyme-substrate complex, rather than via an obligate acyl-enzyme intermediate, with a series of glutamate residues capable of promoting substrate binding and deprotonation in a functionally redundant manner. The extensive acceptor-binding site appears capable of accommodating and stabilising all intermediates formed during the iterative acetylation cycle, but unable to process the *cis*-AMHO monomer or the Fe^3+^–FsC species.

Given that deletion of the *sidG* gene does not impair virulence in *A. fumigatus*^9^, the biological rationale for FsC acetylation warrants some consideration. A clear consequence of acetylation is enhanced chemical stability. TAFC is substantially more resistant to hydrolytic degradation than FsC, particularly under acidic conditions, where the free amino groups of FsC promote intramolecular catalysis of ester cleavage.^26,27^ By neutralising these amino groups, acetylation protects the siderophore scaffold and prolongs extracellular persistence, making TAFC a more robust iron-scavenging molecule. TAFC may also be a better iron chelator than FsC. Although direct thermodynamic comparisons are lacking, chelator-challenge experiments with Ga^3+^ complexes indicate greater metal retention by TAFC than FsC.^28^ Neutralisation of the positively-charged amines of FsC by acetylation may also reduce electrostatic penalties associated with initial binding of Fe^3+^ to form the complex.

Beyond influencing stability and metal coordination, SidG-mediated acetylation of FsC functions as a molecular recognition determinant within the fungal cell. FsC and TAFC utilise distinct uptake transporters (MirD and MirB)^10,11^ as well as different intracellular hydrolases (SidJ and EstB) for degradation.^12–14^ Acetylation therefore directs siderophores into specific acquisition and processing pathways. More broadly, acetylation may also represent a mechanism for siderophore diversification. The ability to produce both FsC and TAFC generates chemically distinct, yet functionally related siderophores, potentially broadening ecological fitness. Such diversification could also increase uptake specificity and reduce susceptibility to siderophore piracy by competing microorganisms. For example, the uptake of the TAFC by *Saccharomyces cerevisiae* via Taf1 highlights the potential for cross-species siderophore exploitation^29^, although whether transporter diversification functions to limit this process remains unknown.

## METHODS

*Molecular cloning and mutagenesis.* The expression construct encoding SidG from *Aspergillus fumigatus* Af293 was obtained by gene synthesis (Epoch Life Sciences) based on the NCBI sequence XM_743592 and subcloned into pET24a containing a custom His₈ tag (see **Supplementary Information** for sequences). Deletions and point mutations to SidG were constructed using the Q5 site-directed mutagenesis kit (NEB), with primers detailed in **Supplementary Table 1**. The resulting PCR products were processed according to the manufactures protocol, and resulting plasmids sequenced to verify the presence of the correct mutation.

*Protein overexpression and purification.* An aliquot of chemically competent *E. coli* BL-21 Star (DE3) cells (50 µL) was transformed with corresponding plasmid DNA. Transformed cells were plated on LB agar supplemented with kanamycin (50 µg / mL) and incubated overnight at 37 °C. A single colony was picked and used to inoculate LB media (10 mL) containing kanamycin (50 µg / mL) and incubated overnight at 37 °C with shaking (180 rpm). The preculture (10 mL) was used to inoculate a flask of LB media (1 L) containing kanamycin (50 µg / mL), and incubated at 37 °C with shaking (180 rpm) until an O.D. of 0.7 – 1.0. Protein production was then induced by addition of IPTG (200 µM) and incubated overnight at 15 °C with shaking (180 rpm). The cell culture was centrifuged (4000 rpm, 15 minutes, 4 °C), and cell pellets resuspended in loading buffer (20 mM imidazole, 20 mM Tris, 100 mM NaCl, 10% glycerol, pH 7.4). Lysing of the cells was carried out by high pressure (20 psi) cell disruption and centrifuged (17,000 rpm, 45 minutes, 4 °C) to pellet insoluble cell debris. The soluble cell lysate was filtered through a 0.45 µm syringe filter and loaded onto a 1 mL HiTrap Ni-NTA FastFlow column. The column was washed with loading buffer (10 mL) followed by washing/elution with increasing concentrations of imidazole (5 mL of: 50 mM; 3 mL of: 100, 200, 300, 500 mM). Protein content of the elution fractions was visualised by SDS-PAGE and desired fractions were concentrated and buffer-exchanged into storage buffer (20 mM Tris, 100 mM NaCl, 10% glycerol, pH 7.4) using a 10,000 MWCO Vivaspin centrifugal concentrator at 4000 rpm. The N-terminal pHis_8_-tag was cleaved by incubation with thrombin overnight at 4 °C. Completion of cleavage was confirmed by ESI-Q-TOF-MS analysis. The resulting cleavage reaction was subjected to size exclusion chromatography on a HiLoad 16/600 Superdex 200 column equilibrated with 20 mM Tris, 100 mM NaCl, 10 % glycerol, pH 7.4 using an ÄKTA Purifier System. Pure fractions, as identified by SDS-PAGE, were concentrated using a 10,000 MWCO Vivaspin centrifugal concentrator. The concentrated protein was aliquoted and flash-frozen in liquid nitrogen and stored at -80 °C.

*Denatured mass spectrometry analysis of intact SidG.* The denatured masses of SidG and variants were obtained using a Bruker MaXis II ESI-Q-TOF-MS connected to a Dionex 3000 RS UHPLC fitted with an ACE C_4_-300 RP column (100 x 2.1 mm, 5 μm, 30 °C). The column was eluted with a linear gradient of 5 – 100 % MeCN containing 0.1 % formic acid over 30 min. The mass spectrometer was operated in positive ion mode with a scan range of 200 – 3000 m/z. Source conditions were: end plate offset at −500 V; capillary at −4500 V; nebulizer gas (N_2_) at 1.8 bar; dry gas (N_2_) at 9.0 L min^−1^; dry temperature at 200 °C. Ion transfer conditions were: ion funnel RF at 400 Vpp; multiple RF at 200 Vpp; quadrupole low mass at 200 *m/z*; collision RF at 2000 Vpp; transfer time at 110.0 µs; pre-pulse storage time at 10.0 µs. Raw mass spectra were deconvoluted and visualised in Bruker Compass DataAnalysis 4.1.

*Native protein mass spectrometry of SidG.* Samples of SidG (190 µM) were buffer-exchanged into 50 mM NH_4_OAc using a 10,000 MWCO Vivaspin centrifugal concentrator (10-fold dilutions, repeated 5 times), and adjusted to 10 µM for analysis. Native mass spectrometry experiments were performed on a Waters SELECT Series Cyclic IMS Mass Spectrometer; a hybrid quadrupole / cyclic ion mobility / time of flight instrument. Samples were infused from a Hamilton syringe (250 uL) using a syringe pump into the ESI source. The mass spectrometer was operated in positive ion mode, with a capillary voltage of 2.5 kV and a scan range of 50 – 8000 *m/z*. Sample-specific settings include: sample cone 30 V, source offset 40–60 V, source temperature 50 °C, desolvation temperature 200 °C, trap collision energy 6 V, trap gas flow 3 mL/min, transfer collision energy 4 V, and transfer gas flow 3 mL/min. Activation of the SidG • acetyl-CoA and SidG • acetyl-CoA • TAFC complexes used quadrupole isolation of the 10^+^ species (*m/z* = 2379.4 and *m/z* = 2456 – 2464, respectively), and an increased transfer collision energy to liberate bound species (50 V and 30 V, respectively). Raw mass spectra were visualised in MassLynx 4.2 and subjected to minimal smoothing.

*In vitro reconstitution of SidG-catalysed acetylation.* SidG (0.5 µM) was incubated with FsC (100 µM) and acetyl-CoA (500 µM) in storage buffer (20 mM Tris-HCl, 100 mM NaCl, pH 7.4) in a total reaction volume of 25 µL at 25 °C. Reactions were quenched at defined time points (10 -600 s) by addition of 50 µL methanol, resulting in protein precipitation. Samples were centrifuged to remove precipitated protein and the resulting supernatants were analysed by UHPLC-HR-ESI-MS. Based on the time-course analysis, the 60 s time point was selected for all subsequent endpoint assays used to compare SidG variants and the 120 s time point was selected for all alternative acyl-CoA substrates. Purified FsC was obtained from *Fusarium roseum* and purified according to previously reported protocols.^30,31^

*Crystallisation and structure determination of SidG (S149A).* The N-terminal His_8_-tag cleaved SidG (S149A) was subjected to size-exclusion chromatography on a HiLoad 16/600 Superdex 200 column equilibrated with 20 mM Tris, 100 mM NaCl, pH 7.4 using an ÄKTA Purifier System. Pure fractions, as identified by SDS-PAGE, were concentrated using a 10,000 MWCO Vivaspin centrifugal concentrator to ∼10 mg/mL and subjected sparse-matrix crystal screening at 20 °C using 96-well sitting drops. Protein crystals were grown at a concentration of ∼10 mg/mL at 20 °C in a 3:2:1 ratio (v/v/v), protein : crystallisation solution (0.1 M MES, 40% (v/v) PEG 400, 5% (w/v) PEG 3000) : seed stock (Molecular Dimensions Nanoseed beads stainless steel kit following manufacturers protocol). Crystals were mounted on appropriately sized mounting loops and flash-frozen for storage in liquid nitrogen without further addition of cryoprotectant. All datasets were collected at the Diamond Light Source synchrotron facility on Beamline I03. Diamonds auto processing pipeline, utilising xia2 with DIALS, was used for indexing, scaling and merging. Molecular replacement and refinements were performed with CCP4i (RefMac5, COOT) and Phenix.^21,32–34^ Data collection, phasing and refinement statistics for SidG(S149A) • acetyl-CoA are reported in **Supplementary Table 2**.

*Oligomeric state determination by size-exclusion chromatography.* Oligomeric state experiments were conducted on an ÄKTA Purifier System using a Superdex 200 HiLoad 16/600 with a flow rate of 1 mL / min. The column was calibrated using Bio-Rad Gel Filtration Standards (Thyroglobulin (670 kDa), γ-globulin (158 kDa), Ovalbumin (44 kDa), Myoglobin (17 kDa), and Vitamin B12 (1.35 kDa)) in 20 mM Tris, 100 mM NaCl, pH 7.4. Elution volumes for each protein standard were used to construct a calibration curve. Following calibration, purified SidG (200 µM, 2 mL) was injected onto the column under the same conditions, and the elution time was used to calculate the molecular weight of the protein using the calibration curve.

*Molecular dynamics simulations of SidG • acetyl-CoA and SidG • acetyl-CoA • FsC complexes.* The SidG • acetyl-CoA and SidG • acetyl-CoA • FsC complexes were modelled with the Protenix server.^24^ For the SidG • acetyl-CoA • FsC complex, the FsC was positioned in two different orientations with respect to acetyl-CoA C=O: Model 0, in which the minimum distance between the FsC amine nitrogen and the thioester carbonyl carbon of acetyl-CoA was 7.0 Å, and Model 3, in which the corresponding distance was 4.8 Å. These were selected as starting structures for the simulations. The positioning of acetyl-CoA was approximately the same and consistent with the crystal structure. In the first instance, a series of three independent repeats of 300 ns cMD simulations were conducted with all amine groups in the NH_3_^+^ form, starting from Protenix Model 0 and Model 3. Subsequently, three independent 400 ns cMD simulations were initiated from frames in which one of the amine groups had been deprotonated by a glutamate residue. The starting geometries were constructed from Frame 9163 from Run 1 of SidG • acetyl-CoA • FsC complex from Protenix Model 3, where the deprotonated amine is hydrogen bonded to both E103 and E141 (**Supplementary Fig. 8a**), and from Frame 10212 from Run 2 of SidG • acetyl-CoA • FsC complex from Protenix Model 0, where the deprotonated amine is hydrogen bonded to both E179 and E141 (**Supplementary Fig. 8b**). Deprotonation by each glutamate residue was considered in turn. Finally, three independent repeats of 400 ns cMD followed by 150 ns aMD were performed on the SidG • acetyl-CoA • MAFC complex. Simulations were initiated from randomly selected frames obtained from the preceding simulations in which the FsC amino group, modelled in its neutral NH_2_ state, adopted a near-attack conformation relative to the acetyl-CoA thioester carbonyl. An acetyl group was then manually appended to the amine to generate the starting structures for subsequent simulations monitoring the evolution of the complex. AMBER trajectories stripped of water and ions can be found as the used custom parameters can be found on Zenodo: https://doi.org/10.5281/zenodo.22754438. A combination of CCPTRAJ^35^, Chimera^36^ and ChimeraX^37^ were used for analysis of the trajectories, ChimeraX was used throughout the course to prepare and visualize the structures.

## DATA AVAILABILITY

The minimum dataset required to interpret, verify and extend the work is provided in the manuscript and Supplementary Information. The structure factor amplitudes and atomic coordinates for SidG • acetyl-CoA have been deposited in the RCSB Protein Data Bank under PDB code 32FB. The protein sequences used in this study are reported in the Supplementary Information. The co-ordinates for the frames shown in **Fig. 5b**, **Fig. 5c**, and **Fig. 6b** along with the cMD trajectories stripped of water and ions with frames every 20 ns are deposited at https://doi.org/10.5281/zenodo.22754439. All materials in this study are available via written request to the corresponding author.

## ACKNOWLEDGEMENTS

This work was supported by a UKRI Future Leaders Fellowship to M.J. (MR/W011247/1), from which Y.T.C.H was funded. J.N. gratefully acknowledges funding from an EPSRC Doctoral Training Partnership (EP/T51794X/1) studentship. F.A. was supported by a UKRI Future Leaders Fellowship (MR/V022334/1). C.D.F. was supported by supported by the interdisciplinary network OI MICROBES and the GS LSH of University Paris-Saclay as part of France 2030 programme ANR-11-IDEX-0003. L.M.A. gratefully acknowledges a Royal Society Research Grant RGS\R2\252466. J.R.L. acknowledges support from EPSRC (EP/Z531200/1, EP/Z535709/1), BBSRC (BB/Z517318/1, UKRI723) and MRC/JPIAMR (UKRI877). The Bruker MaXis II instrument used for intact protein MS analysis was funded by the BBSRC (BB/M017982/1). The Waters SELECT Series Cyclic IMS instrument used for native protein MS work was funded by UKRI (MR/W011247/1). The computing facilities were provided by the Scientific Computing Research Technology Platform at the University of Warwick. The authors are grateful to Prof. Greg Challis for useful discussions relating to iron binding capabilities of FsC and TAFC.

## AUTHOR CONTRIBUTIONS

M.J. and F.A. conceived, designed and supervised the study. J.N. generated *E. coli* expression constructs, performed protein overexpression and purifications, conducted all assays and analysed data. J.N. and M.J. conducted native protein MS analyses. J.N. and Y.T.C.H. performed protein crystallisation and optimisation. J.N., Y.T.C.H., L.M.A. and C.D.F. collected diffraction data and solved the structure of SidG • acetyl-CoA. G.W. provided purified des-ferric fusarinine C. J.R.L. conducted molecular dynamics simulations of SidG and its substrate complexes. M.J. wrote the manuscript with input from all authors.

