## Supplementary Information for "Structural and biochemical characterisation of an iterative GCN5-related *N*-acetyltransferase required for fungal siderophore tailoring"

### 1. Supplementary Figures

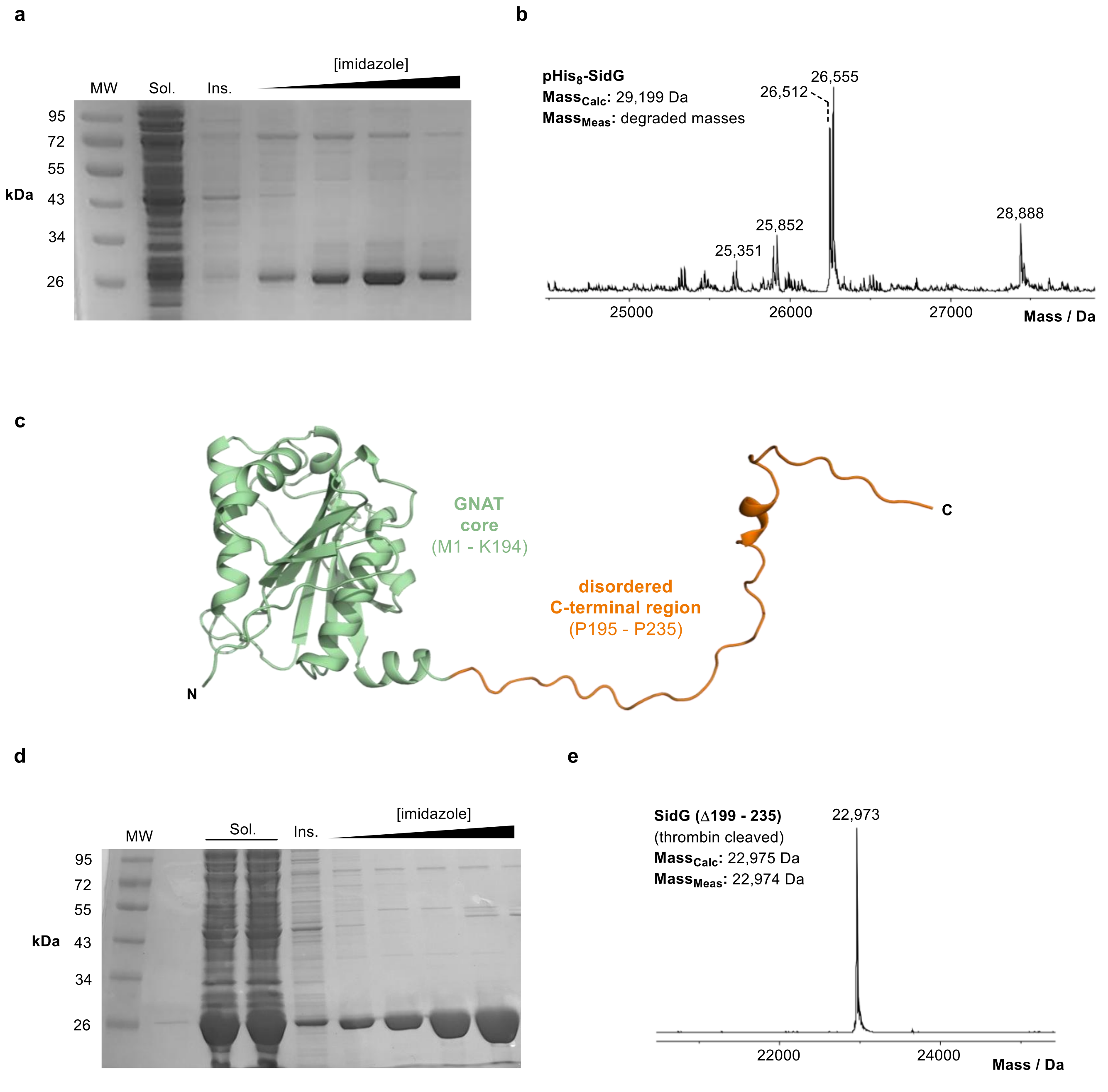

**Supplementary Figure 1. Analysis of recombinant SidG constructs**. (a) 12 % SDS-PAGE gel of full-length pHis$_8$-SidG following IMAC purification. (b) Deconvoluted intact protein mass spectrum of full-length pHis$_8$-SidG. A range of species are observed, all lower than the expected mass, suggesting degradation. (c) AlphaFold model of full-length SidG.[1] The GNAT core region is highlighted in green, and the disordered C-terminus is shown in orange. (d) 12 % SDS-PAGE gel of SidG (Δ199-235) following IMAC purification and thrombin cleavage of the pHis$_8$-tag. (e) Deconvoluted intact protein mass spectrum of pHis$_8$-SidG (Δ199-235).

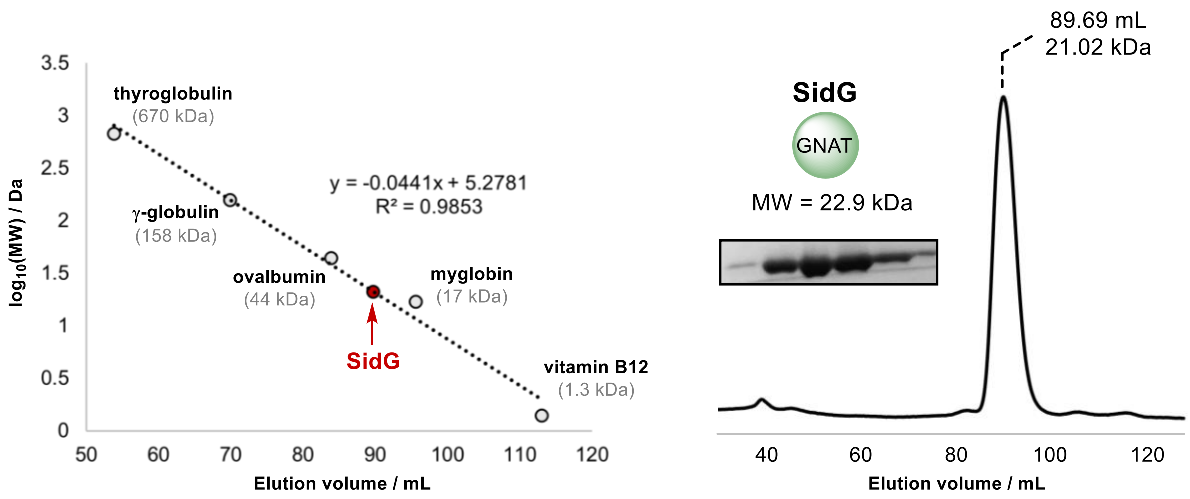

**Supplementary Figure 2. Oligomeric state determination of SidG using size-exclusion chromatography**. Calibration curve obtained from injection of Bio-Rad Gel Filtration Standards. Size-exclusion chromatography profile of SidG yields a single peak corresponding to a monomeric species, with a calculated molecular weight of 21 kDa (actual = 22.9 kDa) and the protein identity confirmed by SDS-PAGE.

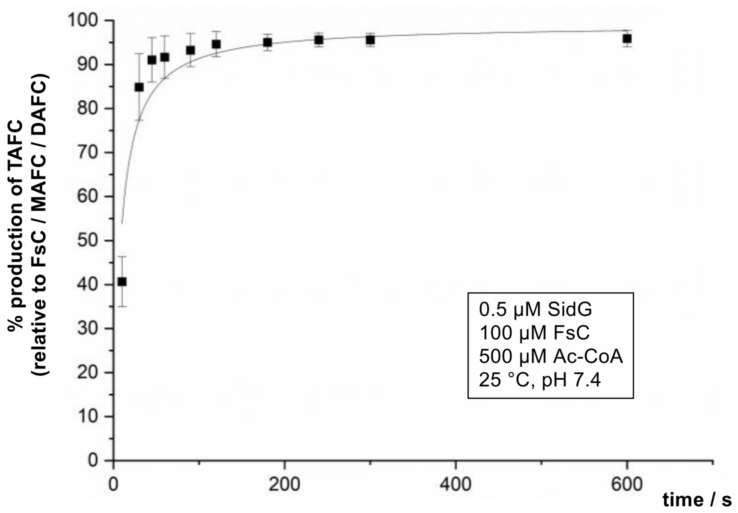

**Supplementary Figure 3. Time course of SidG-catalysed acetylation of FsC.** Plot showing percentage production of TAFC relative to the substrate, FsC, and the intermediates, MAFC and DAFC. The conditions for the time-course assay are shown in the inset. Error bars represent ± 1 standard deviation (1σ) from the mean, where $n$ = 3.

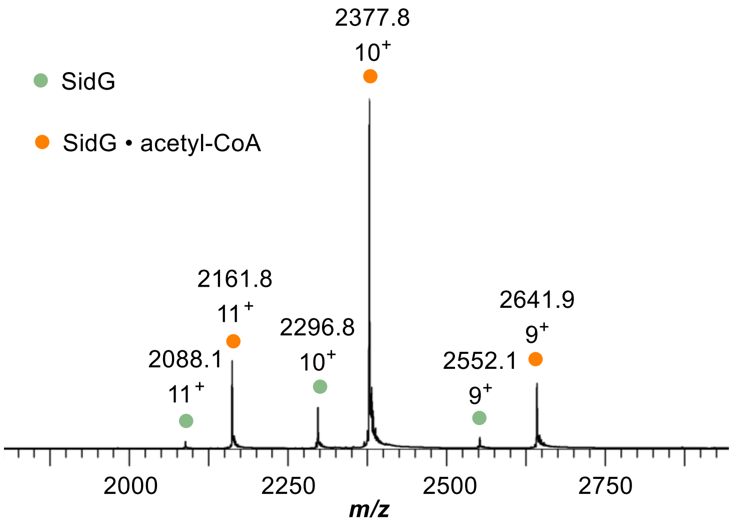

**Supplementary Figure 4. SidG (S149A) purifies with a higher proportion of acetyl-CoA bound.** Native mass spectrum of SidG (S149A) sprayed from 50 mM $NH_4OAc$ showing two charge state distributions from $9^+$ to $11^+$. The green dots indicate the SidG (S149A) protein alone, and the orange dots the SidG (S149A) • acetyl-CoA complex. The amount of acetyl-CoA bound in the S149A mutant is significantly higher than the wild-type (see **Fig. 2a**).

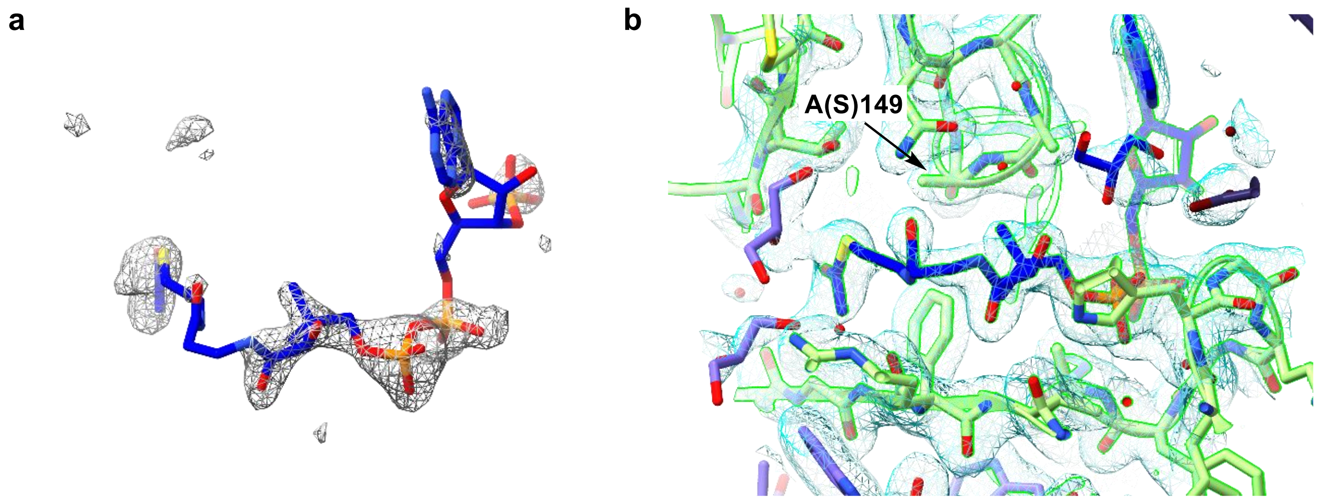

**Supplementary Figure 5. Electron density maps of the acetyl-CoA binding pocket.** (**a**) $Fo{-}Fc$ omit electron density map (brown mesh, contoured at 3.0 $\sigma$) of the acetyl-CoA binding site. (**b**) Final refined structure showing the $2Fo{-}Fc$ electron density map (blue mesh, contoured at 1.0 $\sigma$) covering acetyl-CoA and immediate surrounding residues. The A(S)149 residue is highlighted to illustrate the spatial separation and lack of close contacts with the terminal acetate group of the ligand.

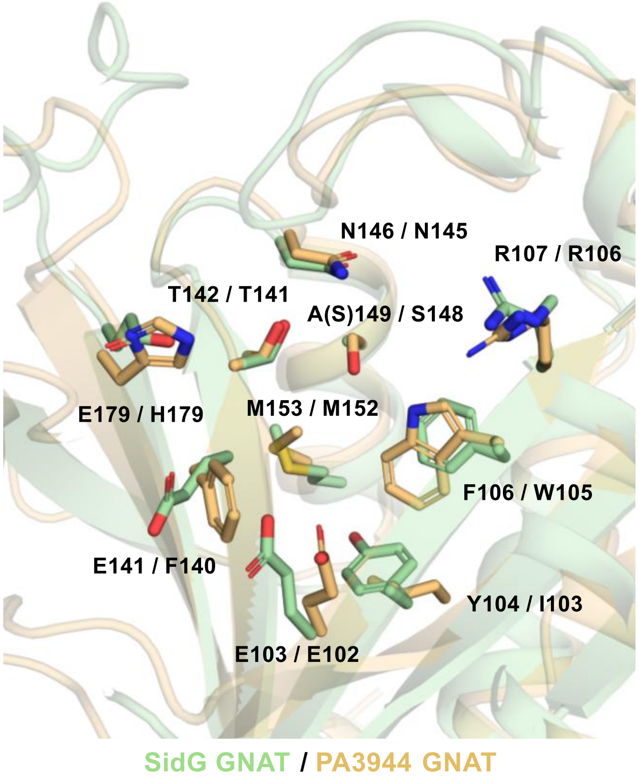

**Supplementary Figure 6. SidG and PA3944 share a conserved GNAT active-site architecture.** Structural overlay of SidG (PDB: 32FB, light green) and PA3944 (PDB: 6EDV, light orange). Residues lining the active site / binding pocket are shown as sticks and labelled according to the respective protein numbering (SidG / PA3944). Core active site residues are conserved between the two GNAT enzymes, including R107, T142, N146 and A(S)149. In contrast, the putative substrate-binding residues E141 and E179 of SidG are replaced by Phe and His in PA3944, respectively, suggesting that these Glu residues represent adaptations for the recognition and binding of FsC.

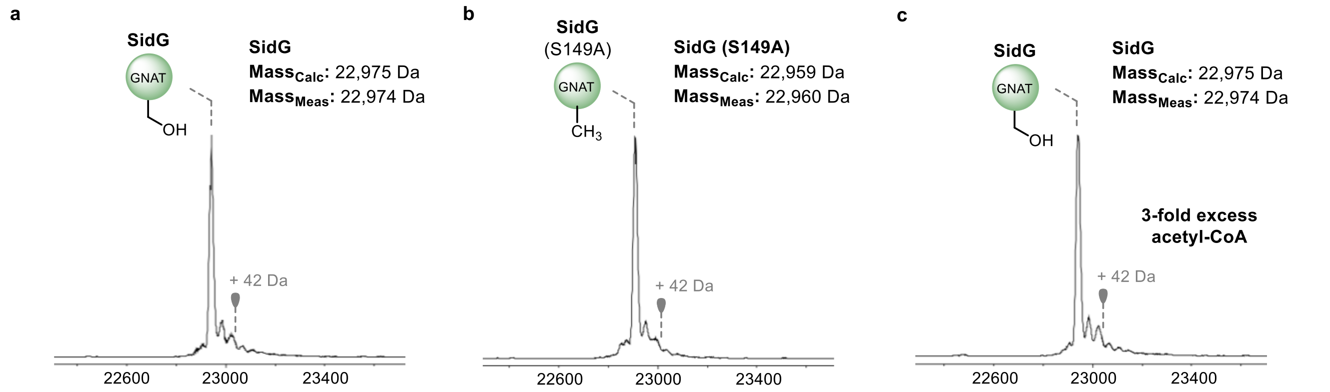

**Supplementary Figure 7. SidG and SidG (S149A) are not acetylated.** Deconvoluted mass spectra of denatured (**a**) wild-type SidG and (**b**) SidG (S149A) following purification. (**c**) Deconvoluted mass spectra of denatured wild-type SidG following incubation with a 3-fold excess of acetyl-CoA. In all cases, the grey marker indicates the position of +42 Da, which would be indicative of acetylation.

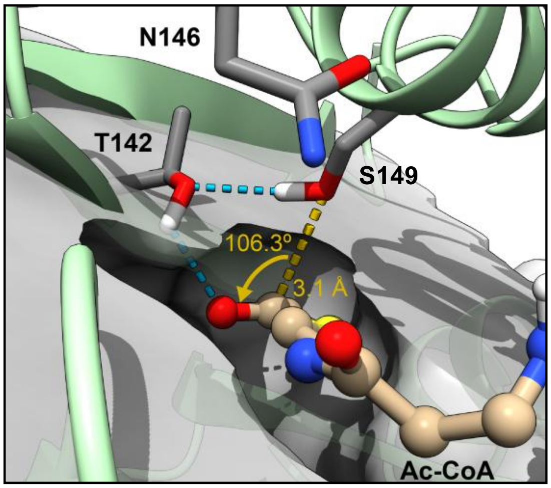

**Supplementary Figure 8. Near-attack conformation for Ser149 with Ac-CoA.** Example frame from a cMD simulation of the SidG • acetyl-CoA complex showing the best observed catalytically competent geometry (Ser149 Oγ→C1 ≤ 3.5 Å; Oγ–C1–O = 100-110°) for acylation of Ser149. The carbonyl oxygen of Ac-CoA does not insert into the putative oxyanion hole formed by T142 and N146 but forms hydrogen bond to T142 side chain. Cyan dashed lines indicate selected hydrogen bonds. Four similar frames were observed in two out of three independent repeats of 300 ns cMD.

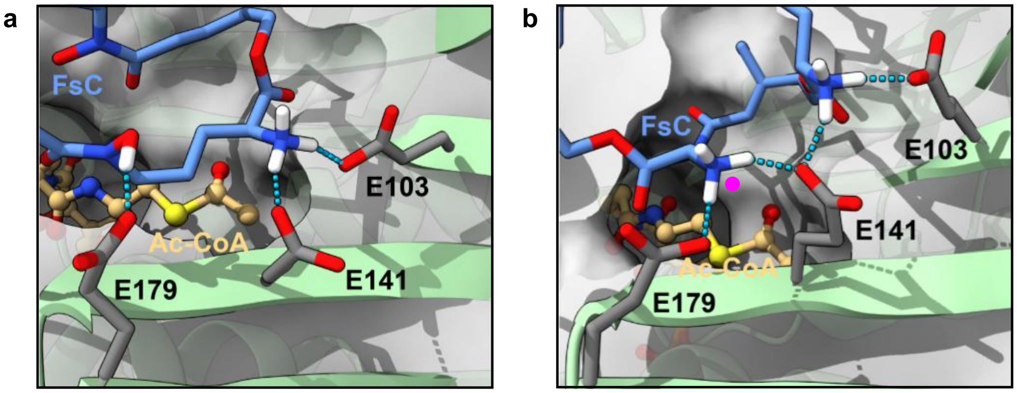

**Protenix:** Model 3 **Protenix:** Model 0

**Supplementary Figure 9. Starting points for SidG • acetyl-CoA • FsC simulations.** Selected frames from 300 ns cMD simulations of SidG • acetyl-CoA • FsC (3 × $NH_3^+$), which were used as starting points for simulations of SidG • acetyl-CoA • FsC (2 × $NH_3^+$, 1 × $NH_3^+$). (**a**) A frame from the simulation starting from Protenix model 3, where the initial distance between the closest amine nitrogen and acetyl-CoA thioester carbonyl was 4.8 Å. Deprotonation of this amine by both E141 and E103 was considered in the subsequent simulations. (**b**) A frame from the simulation starting from Protenix model 0, where the initial distance between the closest amine nitrogen and acetyl-CoA thioester carbonyl was 7 Å. Deprotonation of the amine indicated by a pink dot by both E141 and E179 was considered in the subsequent simulations.

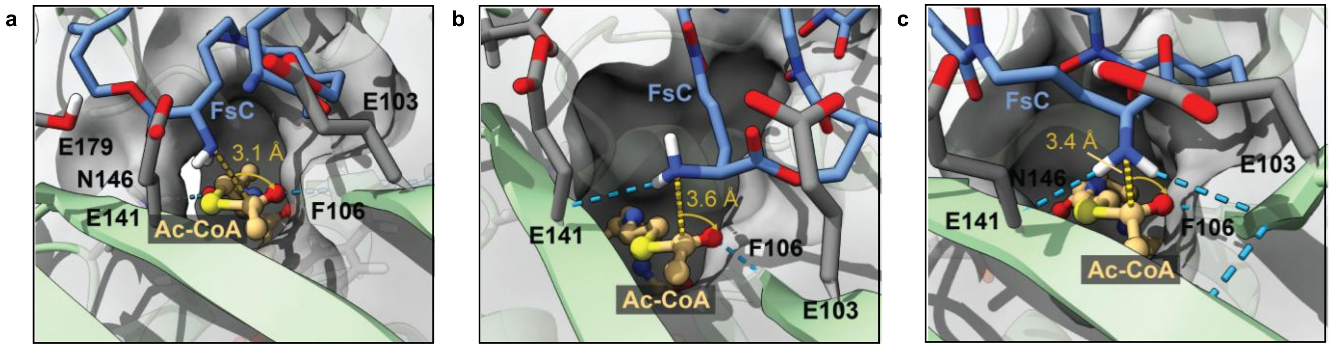

**Starting point:** deprotonation by E179 **Starting point:** deprotonation by E141 **Starting point:** deprotonation by E103

**Supplementary Figure 10. Classical MD simulations of the SidG • acetyl-CoA • FsC ternary complex.** Example frames from simulations of SidG • acetyl-CoA • FsC where one of the amines was deprotonated by either (**a**) E179, (**b**) E141 or (**c**) E103 and where near attack of the FsC amino group ($NH_2$ form) relative to the acetyl-CoA thioester carbonyl was found. The FsC $NH_2$ (N) → acetyl-CoA thioester C=O (C1) distances are shown in the figure. N–C1–O angles were 107.9°, 104.2° and 103.6°, respectively.

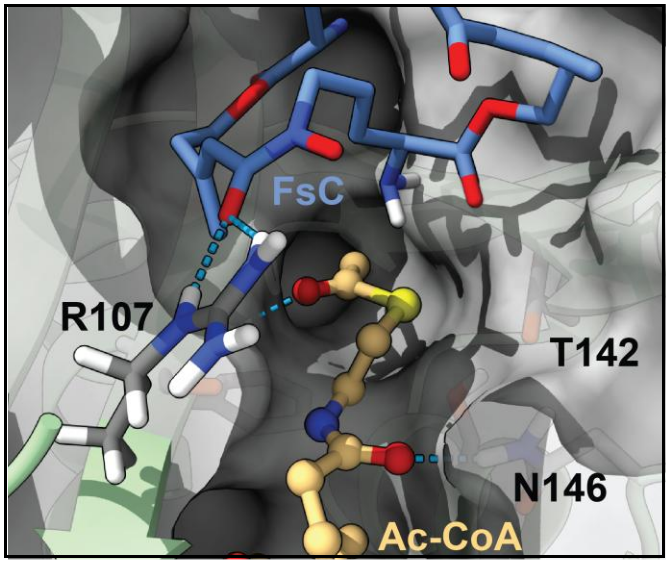

**Supplementary Figure 11. Interaction between R107 and hydroxamate functionality of FsC.** Example frame from cMD simulation of SidG • acetyl-CoA • FsC (same as **Supplementary Figure 10a**) illustrating interaction of R107 with carbonyl of the hydroxamate. Throughout the simulation R107 side chain frequently hydrogen bonds to the oxygens of the hydroxymate moiety facilitating positioning of FsC in the active site.

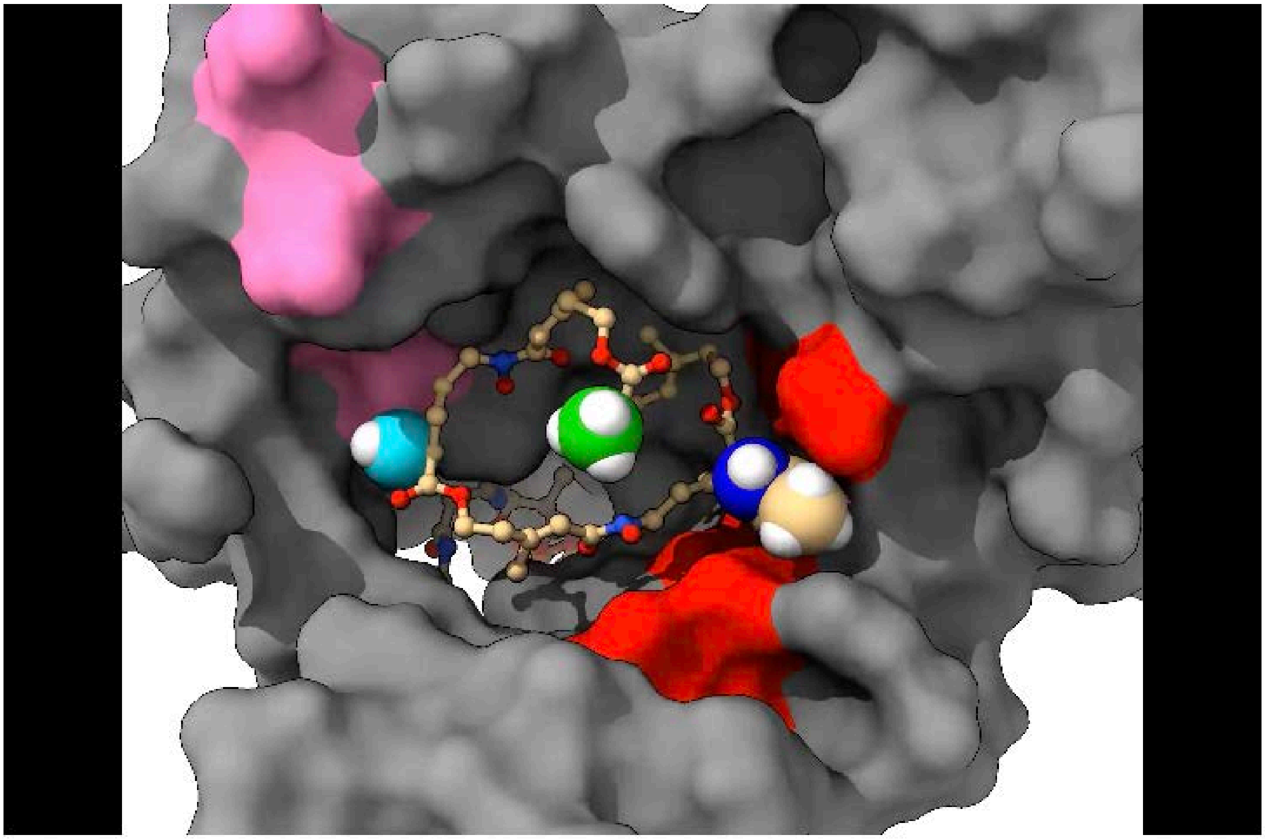

**Supplementary Movie 1. Accelerated MD of SidG • acetyl-CoA • MAFC complex.** Video representation of 150 ns of aMD simulation to monitor the evolution of MAFC in the SidG active site. The starting point of the simulation was constructed by manually attaching an acetyl group to an amine group in a random frame in cMD where a near-attack conformation was found suitable for reaction between acetyl-CoA and the amine of FsC. The acetylated amine is rendered as spheres with the nitrogen coloured in blue. The remaining $NH_3^+$ groups are also rendered as spheres with the nitrogens coloured in cyan and green. The surfaces of E103, E141 and E179 are coloured in red. The surfaces for negatively charged D31, D49, E50 and E92, which transiently interact with NH3+ during the simulation are coloured in pink. The rest of the SidG surface is rendered as a grey surface. At the beginning of the simulation the acetylated amine is trapped between E141 and E103 over the active site. The $NH_3^+$ with the nitrogen coloured in green gets captured by E141 and E103 near the active site around 21 s of the movie (~52 ns into the trajectory). The $NH_3^+$ with the nitrogen coloured in cyan gets captured by E141 and E103 near the active site around 31 s of the movie (~72 ns into the trajectory). Acetyl-CoA, which is present in the active site at the beginning, leaves the binding pocket over the course of this simulation run. In the other two independent repeats, acetyl-CoA remains in the binding pocket but only one of the $NH_3^+$ groups ends up near the active site after the acetylated amine leaves.

### 2. Sequences, Tables and Accession Numbers

#### 2.1. DNA Sequence

```
NCBI Reference Sequence: XM 743592
ATGACAATCAAGGCTCAGCCCACTCTGCACACTGCCCGCCTGGAGCTGGTCCCCCTGGGCCATGAGCATCGCGAGTTCACCAT
GAAGTTGGACATGGACCCCGAGGTCATGAAGATGGTCGCCTTCGGCCGGCCATTTACCGAAGACGAAGCAATCCAGGTTCATA
CCTGGCTGATGAACTGCGCAACGTCGGTGCCTGGCTTCGGAACCTGGGTCGGCTTTGCCGAAGGCGAGTTCGTGGGTTGGTGG
GTATTGGCTCCCGTCCCCACGACGGAGAACCCCAAGAGCTTCAGGACCGATCGAACGGAGTATGGCTTCCGAGTCTCGCCGAA
GTTCTGGGGCCAAGGGTACGCGAAGGAGGGGGCTCGGGAGATGGTTCGCTATGCCTTCGAGGAACTGGGTCTGGCCGAGGTGA
TTGGCGAGACGATGACTATCAACATGGCTTCGCGGGCGGTGATGGCCGGGTGCGGGTTGACGCACGTTGAGACCTTCTTCAAC
AAGTACGATACTCCACCGCCAGGCATTGAGGAGGGAGAGGTACGGTATTCGATCACTAGGGAGGAATGGTTGCGGATGCAAAA
GCCCAGCATGACTCGAAGTCGCTGGTTTCCGGCTTTCGCCAGCTGGCTGCCCCGATTGCTCCTGTCCAGGCTCTGGTCCTATA
TTTTCCAAGGGCGTAGGCTCGCAGCTGGGGCGGCCAGCCCCTGA
```

#### 2.2. Amino Acid Sequences

**pHis$_8$-SidG**

---

```
MKHHHHHHHHGGLVPRGSHGS-

        10         20         30         40         50         60
MTIKAQPTLH TARLELVPLG HEHREFTMKL DMDPEVMKMV AFGRPFTEDE AIQVHTWLMN

        70         80         90        100        110        120
CATSVPGFGT WVGFAEGEFV GWWVLAPVPT TENPKSFRTD RTEYGFRVSP KFWGQGYAKE

       130        140        150        160        170        180
GAREMVRYAF EELGLAEVIG ETMTINMASR AVMAGCGLTH VETFFNKYDT PPPGIEEGEV

       190        200        210        220        230
RYSITREEWL RMQKPSMTRS RWFPAFASWL PRLLLSRLWS YIFQGRRLAA GAASP
```

**pHis$_8$-SidG (Δ199 – 235)**

---

```
MKHHHHHHHHGGLVPRGSHGS-

        10         20         30         40         50         60
MTIKAQPTLH TARLELVPLG HEHREFTMKL DMDPEVMKMV AFGRPFTEDE AIQVHTWLMN

        70         80         90        100        110        120
CATSVPGFGT WVGFAEGEFV GWWVLAPVPT TENPKSFRTD RTEYGFRVSP KFWGQGYAKE

       130        140        150        160        170        180
GAREMVRYAF EELGLAEVIG ETMTINMASR AVMAGCGLTH VETFFNKYDT PPPGIEEGEV

       190
RYSITREEWL RMQKPSMT
```

### 2.3. Supplementary Tables

**Supplementary Table 1.** Primers used for SidG mutagenesis, with associated annealing temperatures.

| SidG Mutant | Primers For / Rev (5'→3') | Annealing Temperature (ºC) |
|---|---|---|
| Δ199-235 | FWD: CAGCATGACTTGAAGTCGCTGGT<br>REV: GGCTTTTGCATCCGCAAC | 66 |
| S149A | FWD: CAACATGGCTGCGCGGGCGGTGA<br>REV: ATAGTCATCGTCTCGCCAATCACCTCGG | 72 |
| T142V | FWD: GATTGGCGAGGTGATGACTATCAACATGGCTTCG<br>REV: ACCTCGGCCAGACCCAGT | 68 |
| N146L | FWD: GATGACTATCCTGATGGCTTCGCGGG<br>REV: GTCTCGCCAATCACCTCG | 64 |
| T142V, N146L* | FWD: GATTGGCGAGGTGATGACTATCCTGATGGCTTC<br>REV: ACCTCGGCCAGACCCAG | 68 |
| R107Q | FWD: GTATGGCTTCCAGGTCTCGCCGAAGTT<br>REV: TCCGTTCGATCGGTCCT | 66 |
| E103L | FWD: CGATCGAACGCTGTATGGCTTCC<br>REV: GTCCTGAAGCTCTTGGG | 60 |
| E141L | FWD: GGTGATTGGCCTGACGATGACTATCAAC<br>REV: TCGGCCAGACCCAGT | 63 |
| E179L | FWD: TGAGGAGGGACTGGTACGGTATTCGATCAC<br>REV: ATGCCTGGCGGTGGA | 64 |

* the plasmid for the SidG N146L mutant was used as the template to introduce the T142V mutation.

**Supplementary Table 2.** Data collection, phasing and refinement statistics for SidG(S149A) • acetyl-CoA.

| **Data collection** | |
|---|---|
| X-ray source | DLS, I03 |
| Temperature (K) | 100 |
| Wavelength (Å) | 0.976254 |
| Space group | C121 |
| Cell dimensions: | |
| a, b, c (Å) | 123.88, 48.98, 35.97 |
| α, β, γ (˚) | 90.00, 106.46, 90.00 |
| Monomers per asym. unit | 1 |
| Resolution (Å) | 45.28-2.10 (2.13-2.10) |
| Rmeas | 0.1058 (0.8065) |
| <I/σ(I)> | 13.04 (1.12) |
| CC1/2 | 0.9979 (0.7767) |
| Wilson B-factor ($Å^2$) | 37.18 |
| Total no. reflections | 83718 (20523) |
| No. unique reflections | 12228 (3007) |
| Completeness (%) | 99.90 (99.64) |
| Redundancy | 6.85 (6.75) |
| **Refinement** | |
| Resolution (Å) | 34.5 - 2.098 (2.31- 2.10) |
| No. unique reflections | 12224 (3005) |
| Rwork / Rfree | 0.1838/0.2367 (0.2653/0.2805) |
| No. atoms | 1535 |
| Protein | 1419 |
| Ligand | 69 |
| Solvent | 47 |
| Average B-factor ($Å^2$) | 46.3 |
| Protein | 45.46 |
| Ligand | 60.78 |
| Solvent | 50.31 |
| R.M.S. deviations | |
| Bond length (Å) | 0.67 |
| Bond angles (Å) | 1.05 |
| Ramachandran | |
| Favoured (%) | 97.71 |
| Allowed (%) | 2.29 |
| MolProbity | |
| Clashscore | 5.55 |
| MolProbity score | 1.71 |
| PDB code | 32FB |

**3. Additional Experimental Details**

Molecular dynamics (MD) simulations were performed using the AMBER ff19SB force field.[2] All the ligands were parametrised using ANTECHAMBER[3] and GAFF2 force field with abcg2 charge model.[4] The charge states of the amino acids at pH 7.4 were evaluated using the H++ webserver[5]or ProteinPrepare.[6] The resulting structure was neutralised with $Na^+$ ions and solvated with OPC water[7], such that no atom belonging to the complex was less than 10 Å from any box edge, using the LEaP module.[8] Additional $Na^+$ and $Cl^-$ ions were added to obtain 100 mM final salt concentration. MD heating, equilibration, and production steps were performed using the combination of CPU and GPU accelerated AMBER software on an HPC cluster equipped with NVIDIA RTX6000 or L40 graphics cards. Simulation used the SHAKE algorithm[9] to constrain all protein bonds involving a hydrogen atom; a 2.0 fs time-step was used in these simulations. Long-range electrostatics were calculated using the Particle Mesh Ewald (PME) method with a 12.0 Å cut-off.[10] PME was used for nonbonded interactions. In all simulations, the Langevin thermostat ($\gamma$ = 2.0 $ps^{-1}$) was used to maintain temperature control.[11] The solvated protein was then equilibrated by carrying out a short minimization, 50 ps of heating and 50 ps of density equilibration with weak restraints on the protein followed by 500 ps of constant pressure equilibration at 300 K. After a two-step minimization process, in which solvent molecules were allowed to relax before the entire system was minimized, the system was slowly heated to 300 K over 0.1 ns in a canonical ensemble (NVT) simulation, then equilibrated for 2 ns by performing isothermal-isobaric (NPT) simulations at 300 K using a Berendsen barostat.[12] For each system 3 independent repeats of 300 ns isothermal-isobaric (NPT) production simulations using classic approach (with no acceleration) were performed using a Monte-Carlo barostat[13] with simulation frames written every 20 ps for analysis. For all simulations, long-range electrostatics were calculated using the PME method with an 8.0 Å cutoff.

For SidG • acetyl-CoA • FsC complex with 1 x $NH_2$ + 2 x $NH_3^+$ additional 300 ns of accelerated molecular dynamics was performed from the restart points of classical MD runs. The aMD modification of the potential was defined as:

$$\mathrm{V(r)^* = V(r) + \Delta V(r)}$$

$$\mathrm{\Delta V(r) = \frac{(E_P - V(r))^2}{(\alpha_P + E_P - V(r)} + \frac{(E_D - V(r))^2}{(\alpha_D + E_D - V_D(r)},}$$

where V(r) is the normal potential and $V_D(r)$ is the normal torsion potential, with $E_P$ = -122623.99 to -122657.6027 kcal $mol^{-1}$; $a_P$ = 8362.6 $mol^{-1}$; $E_D$ = 1824.8748 to 1842.1956 kcal $mol^{-1}$; $a_D$ = 133 kcal $mol^{-1}$ (estimated from the cMD simulation, see Amber manual).

A combination of CCPTRAJ[14], Chimera[15] and ChimeraX[16] were used for analysis of the trajectories, ChimeraX was used throughout the course to prepare and visualize the structures.

---
